# Asymmetric evolution of the transcription profiles and *cis*-regulatory sites contributes to the retention of transcription factor duplicates

**DOI:** 10.1101/115857

**Authors:** Nicholas L. Panchy, Christina B. Azodi, Eamon F. Winship, Ronan C. O’Malley, Shin-Han Shiu

## Abstract

Transcription factors (TFs) play a key role in regulating plant development and response to environmental stimuli. While most genes revert to single copy after a duplication event, transcription factors are retained at a significantly higher rate. However, it is unclear why TF duplicates have higher rates of retention relative to other genes. In this study, we compared three types of features (expression, sequence, and conservation) of retained TFs following whole genome duplication (WGD) events to genes with other functions, using *Arabidopsis thaliana* as a model. We found that gene function groups with higher maximum expression but lower mean expression tended to have higher duplicate retention rate post WGD, though TFs in particular are retained more often than would be expected based on the features examined. Conversely, expression of individual genes was not associated with duplication, but sequence conservation was. Furthermore, we found that the evolution of TF expression patterns and cis-regulatory cites favors the partitioning of ancestral states among the resulting duplicates. In particular, we found that one duplicate retains the majority of ancestral expression and cis-regulatory sites, while the “non-ancestral” duplicate was enriched for novel regulatory sites. To investigate how this pattern of partitioning pattern evolved, we modeled the retention of ancestral states in duplicate pairs using a system of differential equations. Our findings indicate that duplicate pairs evolve to a partitioned state more often than away from it, which in combination with accumulation of new regulatory sites in non-ancestral duplicates, suggest that selection favors partitioning via neofunctionalization.

**Author Summary:** Gene expression is controlled by regulatory proteins known as transcription factors. These factors control how an organism develops and responds to its environment. The evolution of transcription factor functions also contributes to the emergence of new species and crop domestication. In plants, new transcription factors mainly arise due to polyploidy, multiplication of the genome. Although most duplicated copies are lost following a genome duplication event, transcription factors are exceptional because they are often kept. Furthermore, we found that transcription factor duplicates that tend to diverge in how they are expressed and regulated in an unusual way where one copy mirrors the original, pre-duplication functional states of the ancestral gene, while the other loses the ancestral status and instead accumulates novel regulatory sites. Our results suggest these duplicate transcription factors may have been kept because one copy preserve ancestral function while the other has evolved new ones.

## Introduction

Plant genomes are replete with paralogous genes derived from a variety of duplication events and mechanisms (Panchy et al., 2016). Among them, whole genome duplication (WGD) events are responsible for the majority of extant duplicate genes (Panchy et al. 2016). Analysis of sequenced plant genomes has revealed evidence for two ancient WGD events prior to the divergence of angiosperms (Jiao et al. 2011) and, since then, more than a dozen WGD events have occurred across a variety of angiosperm lineages (Lyons et al. 2008; Lee et al. 2013; Myburg et al. 2014; Renny-Byfield et al. 2014; Soltis et al. 2014; Wang et al. 2014), including three in the lineage leading to *Arabidopsis thaliana* (Bowers et al. 2003). This suggests that WGD occurs more frequently in plants relative to other lineages, as the last known WGD event in the *Saccharomyces cerevisiae* (Wolfe and Shields 1997; Kellis et al. 2004) and human (Panopoulou et al. 2003; Dehal and Boore 2005) lineages occurred prior to the radiation of angiosperms. Due to frequent WGD events and ensuing gene losses, the number of genes in extant angiosperms vary widely but certain gene families, particularly transcription factor (TF) families, have expanded dramatically in plant genomes (Lespinet et al. 2002; Shiu et al. 2005).

The duplication of TFs is of particular interest because duplicate TFs contribute significantly to plant adaption (Lehti-Shiu et al. 2016), agricultural traits (Zhang et al. 2011), and domestication (Liu et al. 2015). Not only does the expansion of several TF families coincide with major events in the evolution of plants (i.e. migration to land and expansion of flowering plants,Weirauch and Hughes 2011), but TF duplication has specifically influenced the evolution of flowering time (Schranz et al. 2002), floral structures (Theissen and Melzer 2007) and the co-option of floral regulators to fruit development (Litt and Irish 2003, McCarthy et al. 2015). WGD accounts for ~90% of the expansion of TF families across plants lineages (Maere et al. 2005). Compared to other eukaryotes, plant TF duplicates derived from WGD are retained at higher rates than most plant genes with other functions (Seoighe and Gehring 2004; Shiu et al. 2005).

Furthermore, TFs are consistently enriched among WGD duplicates across divergent plant species (Carretero-Paulet and Fares 2012), although the overall retention rates in these species lineages vary greatly (Lynch and Conery 2000; Lynch and Conery 2003; Moghe and Shiu 2014). Because WGD results in duplication of all genes in a genome, the observed differences in the expansion of different gene families (e.g. Blanc and Wolfe 2004; Seoighe and Gehring 2004; Hanada et al. 2008; Li et al. 2016) must result from differential rates of gene retention. Previously, sequence features (e.g. gene length), biochemical activities (e.g. expression level), evolutionary characteristics (e.g. substitution rates), and annotated function have been used to assess the general properties of retained duplicates in plant genomes (Jiang et al. 2013; Moghe et al. 2014). These properties of extant TFs, however, tell us little about the evolutionary trajectories of TF duplicates. In order to determine what properties are associated with retained TFs and how well those properties explain the differences in retention rates between TFs and other function groups, it is critical to include information about how retained pairs have changed since duplication.

Modeling approaches to infer ancestral expression levels based on extant gene properties have been used to determine the rate of gene activation and repression in duplicate genes in *Drosophila melanogaster* (Oakley et al. 2006) and analyze the evolution of stress response in *A. thaliana* (Zou et al. 2009b). This approach allows for the explicit characterization of how duplicate TFs may have deviated (or not) from their ancestral state over the course of evolution, which in turn may provide information about the mechanism(s) contributing to TF retention. Among theories proposed to explain how duplicated genes may be preserved, subfunctionalization (Force et al. 1999) and gene balance (Birchler and Veitia 2007; Birchler and Veitia 2010; Baker et al. 2013) have been hypothesized to explain the retention of TFs in particular, as well as other genes with high interactivity (Seoighe and Gehring 2004; Maere et al. 2005; Shiu et al. 2005). In addition, neo-functionalization (Ohno, 1970) and escape from adaptive conflict (Des Marais and Rausher 2008) can lead to retention of both duplicate copies through selection on new function or novel optimization of existing functions respectively. Nevertheless, to assess the contribution of these different mechanisms, knowledge of the ancestral functional states of a TF duplicate is essential.

In this study, we compared the retention rate of TF WGD duplicates against all other genes in *A. thaliana* and against genes in each of 19 other “function groups” of similar size to TFs. First, to understand the differences in retention rates amongst different function groups of genes, we modeled retention rate as a function of the expression, structural features, and conservation genes in each function group. Using the same feature set, we also classified genes within each function group as either duplicated or not. In particular, we were interested to see what features of duplicate genes distinguish them from non-duplicates. In addition, we examined whether the correlations between a feature and retention status was consistent across function groups or if some groups, like TFs, deviated from the norm. Second, we asked to what degree the ancestral and extant functions of duplicate pairs were related to their retention or loss. To determine how gene expression and cis-regulatory sites of TF duplicates have likely evolved post WGD we inferred the ancestral expression and cis-regulatory states of existing TF duplicates. We then asked whether the evolution of TF functions as defined by cis-regulatory sites and expression patterns were correlated, which would be expected if regulatory sites are controlling expression. Finally, to test if natural selection is contributing to the observed distribution of ancestral states amongst extant duplicates, we modeled the evolution of TF WGD duplicates as a system of differential equations which tracks the change in frequency of duplicate pairs retaining the ancestral state in both, one, or neither.

## Results

### Retention rates of duplicate genes in different function groups following WGD

To assess the factors contributing to the differential retention of TF duplicates from WGD events and duplicates from WGD events involved in other functions, we first quantified the retention rates of *A. thaliana* WGD duplicates in 20 different function groups. These function groups include TFs annotated by the Plant Transcription Factor Database (Jin et al. 2014) and 19 other groups defined based on Gene Ontology (GO, Ashburner et al. 2000) molecular functions (see **Methods, Table S1**). Within each function group, genes were classified as “WGD-duplicates” (both duplicate copies retained) or “WGD-singletons” (only one copy retained) depending on whether there were paralogs in corresponding duplicate blocks (Bowers et al. 2003). Because duplicate retention rates are expected to differ across different WGD events, duplicate pairs derived from the α, β, and γ WGD events (Bowers et al. 2003) were analyzed separately. To represent the duplicate retention rate of genes in a function group, we calculated the log odds ratio of genes having a retained WGD-duplicate derived from a specific WGD event for each function group relative to all *A. thaliana* genes (see **Methods**). Thus a more positive retention rate indicates that a function group contains a proportionally higher number of retained WGD-duplicates compared to the genome average, while a more negative rate indicates a proportionally lower number of duplicates. Among the 20 function groups examined, the retention rates were highly heterogeneous and only TFs and protein kinases had retention rate significantly higher than the genome average for all three WGD events (Fig 1).

**Figure 1.**
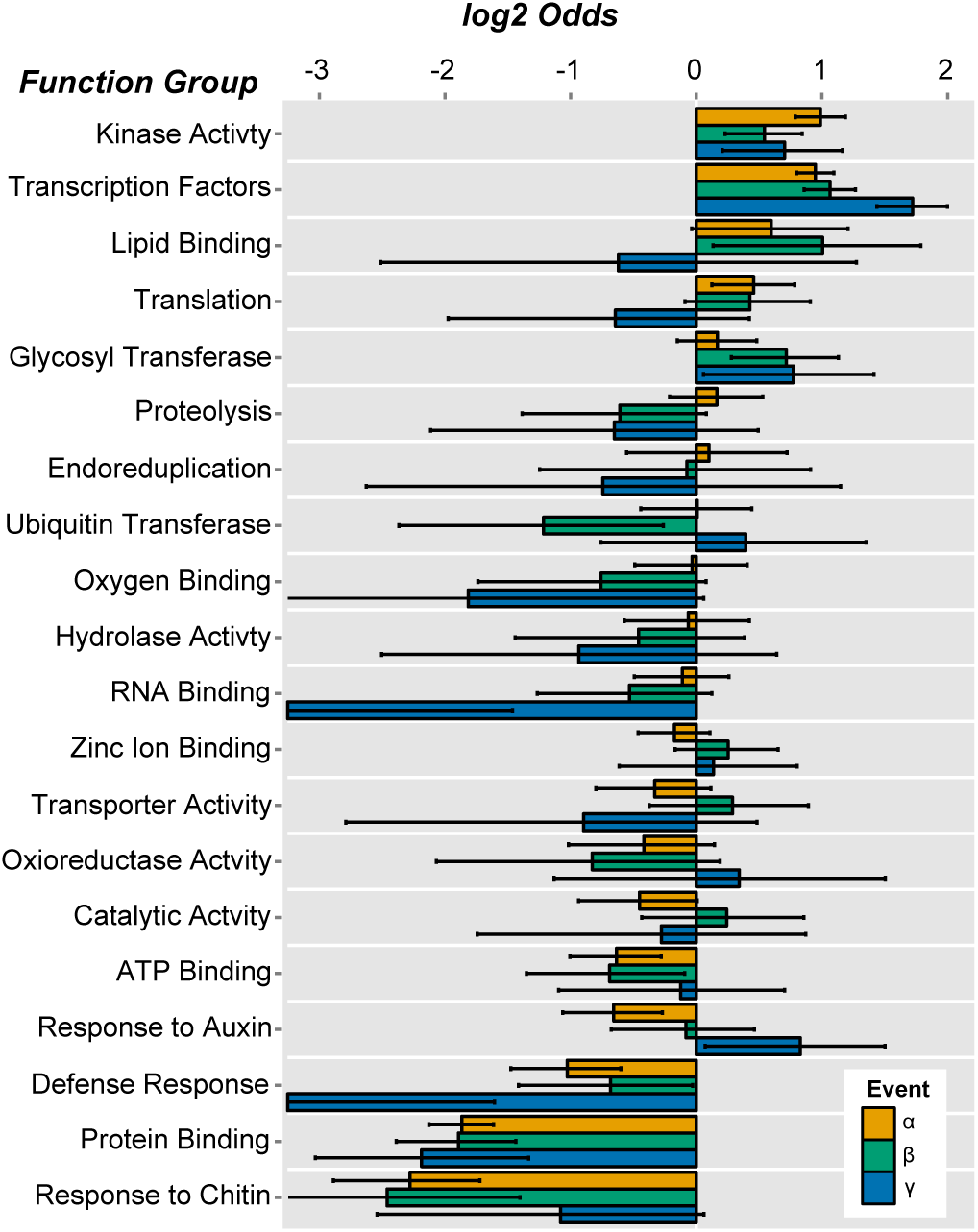
Retention of WGD-duplicate genes in *A. thaliana* The odds of duplicate gene retention within 20 function groups relative to whole genome. Groups are ordered by the odds of duplicate retention in the alpha event. Colors represent different WGD duplication events (α = orange, β = green, γ = blue). Error bars indicate the 95% confidence interval of the odds of retention as determined by the Fisher’s Exact Test implemented in R.

Compared to protein kinases, the retention rate of TFs are even higher for the older (β and γ) duplication events, indicating that on average, the longevity of TF WGD-duplicates is higher than that of protein kinases. Furthermore, the retention rates for the γ TF duplicates (log2(odds) 1.72) is significantly higher than α (log2(odds) 0.95) and β (log2(odds) 1.07) TF WGD-duplicates. This is in spite of the fact that the γ WGD occurred prior to the divergence of monocots and dicots, making it at least twice as old as the α event (Bowers et al. 2003). Importantly, the higher rate of retention for TF WGD-duplicates from the γ event cannot be explained by the fact that these WGD-duplicates went through subsequent duplication events (i.e. α and β). Rather, either more TFs were retained immediately following the γ WGD (i.e. before subsequent duplication) than either the α or β event, or WGD-duplicate retention in subsequent WGD events favored genes with retained WGD-duplicates following the γ WGD (see Supplementary File S1). In summary, TFs were retained more frequently post WGD than most other function groups. Additionally, the retention rate of γ WGD-duplicate TFs is significantly higher than other events, potentially reflecting the contribution of γ WGD-duplicate TFs to the early radiation of flowering plant species.

### Linear model of WGD-duplicate retention rates across function groups

Amongst function groups, TFs stand out as being one of only two which are retained more often than the genome average consistently across all WGD events (the other being protein kinases, one of the largest gene families in plants, Lehti-Shiu and Shiu, 2012). For the rest of the function groups, the retention rate varies above and below the genome average across WGD events without any clear association to either more recent or ancient WGDs. Additionally, although a few highly retained functions also have a higher gene count, there is no clear relationship between the size of a function group and its retention rate for any WGD event (Fig 2A). Therefore, the reason for retention rates differences remains unclear but must involve factors beyond gene function and timing of duplication. To address why retention rates differ, we examined sequence, expression, conservation, and interactivity related features (Fig 2B, **Table S2**) of WGD-duplicate and WGD-singleton genes. We also asked how well the retention rate differences between function groups can be explained by these different features.

**Figure 2.**
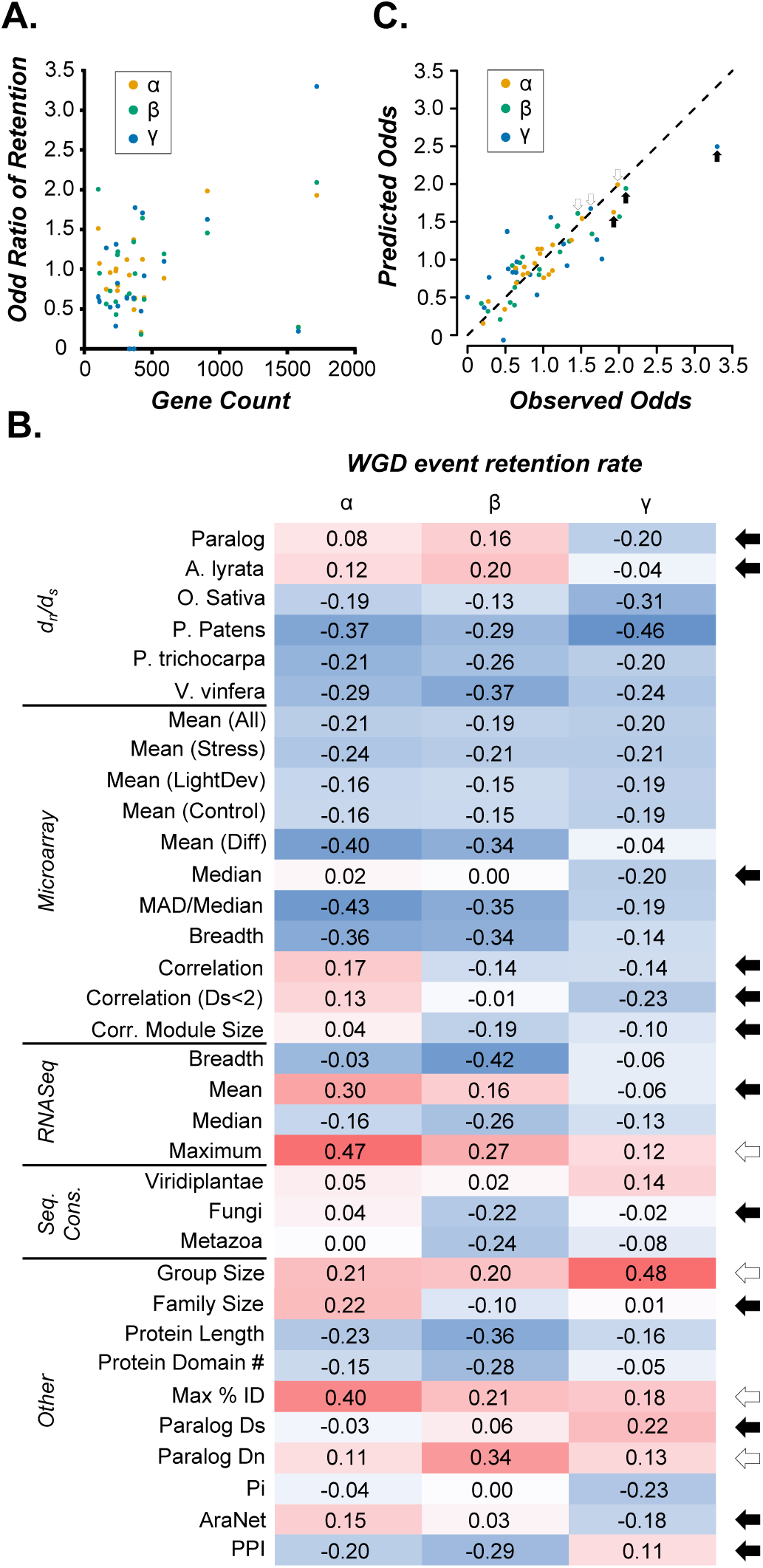
**(A)** Relationships between gene counts and retention rate of WGD duplicates across functional groups (α = orange, β = green, γ = blue). (B) A heatmap of the Pearson product-moment correlation coefficient between the values of a feature across different function groups (y-axis) and the retention rate of functions groups from a particular WGD event (x-axis, indicated by the symbols α, β, and γ). Darker red: stronger positive correlation. Darker blue: stronger negative correlation. Features with different sign of correlation across WGD events are indicated by black arrows. Features with a large (~0.20) difference in magnitude with the same sign are indicated by white arrows. **(C)** The observed odds of duplicate retention (x-axis) for each group plotted against the predicted odds of retention (y-axis) from the best model for each event (α = orange, β = green, γ = blue). The dotted line represents equality between predicted and observed odds of retention. Values from TFs are indicated by a black arrow while values from kinases are indicated by a white arrow.

To see how well the features we considered could explain the retention rate differences among function groups, we constructed a linear model of retention rates for each WGD event. A subset of the features examined here has previously been shown to be significantly associated with retention of WGD-duplicates as a whole (Jiang et al. 2013), however we choose to separate events in this case both because there was a large variance in retention across events within our function groups (Fig 1). Furthermore, WGD events were separated because the correlations between retention rates and feature values have different signs (see black arrows, Fig 2B) and magnitudes (see white arrows, Fig 2B) depending on the WGD events. Hence, a model which describes the relationship between the average features of function groups and duplicate retention rates for a particular WGD event may not be generalizable to other events. Beginning with the full set of 34 features, for each WGD event we determined the subset of features (between 5 and 6 in each case) which maximized the F-statistic of the model (see **Methods**, Table 1). Full model equations can be found in **Supplementary File S2**. Our models explained 87%, 83%, and 65% of the variance in retention rates for the α, β and γ events respectively. Applying the F-test to the maximum F-statistic for each model, we found that each model performs significantly better at explaining the retention rates of function groups than the null model (i.e. fitting retention rates to their mean) at p-value threshold of 0.05.

**Table 1.** Statistics for the best fitting model of duplicate retention rate for each WGD-event.

| WGD Event | # Features<sup>1</sup> | CoD<sup>2</sup> | F-statistic<sup>3</sup> | p-value<sup>4</sup> |
| --- | --- | --- | --- | --- |
| $\alpha$ | 6 | 0.87 | 13.8 | 5.6E-05 |
| $\beta$ | 5 | 0.83 | 13.2 | 7.1E-05 |
| $\gamma$ | 5 | 0.65 | 5.1 | 7.2E-03 |

1: The number of explanatory variables (features) used in the best fitting model
2: Coefficient of Determination ( $R^2$ )
3: the F-statistic is a measure of the goodness of fit of the model to the observed retention rate.
4: p-value of fit based on the F-statistic. A significant p-value ( $p < 0.05$ ) indicates that the model performs better than fitting the mean value to the data, after accounting for the number of features in the model.

To determine the importance of individual features in explaining the retention rate differences among function groups, we determined the change in explained variance caused by independently removing each feature from the model (Table 2). As expected, features which are used in more models have a greater impact on variance explained when removed. Note, maximum expression (RNA-seq) with a positive sign and expression mean (AtGenExpress microarray) with a negative sign were important to models for all three WGD events. This would suggest that more specific expression (i.e. lower average across all conditions, but higher maximum expression under a few specific conditions) increases the likelihood of duplicate retention. Additionally, fewer domains, lower expression correlation with paralogs, and lower nucleotide diversity, were all significantly features of the β and γ models, indicating that these features had a greater impact on older duplication events. These findings suggest long term retention of duplicates favors genes experiencing stronger purifying selection at the primary sequence level (low nucleotide diversity) than at the level of gene expression patterns (lower expression correlation reflecting higher degrees of expression divergence). The remaining feature were found in only one of the models and had significant but much smaller impacts on the variance explained than the features discussed above (Table 2).

**Table 2.**
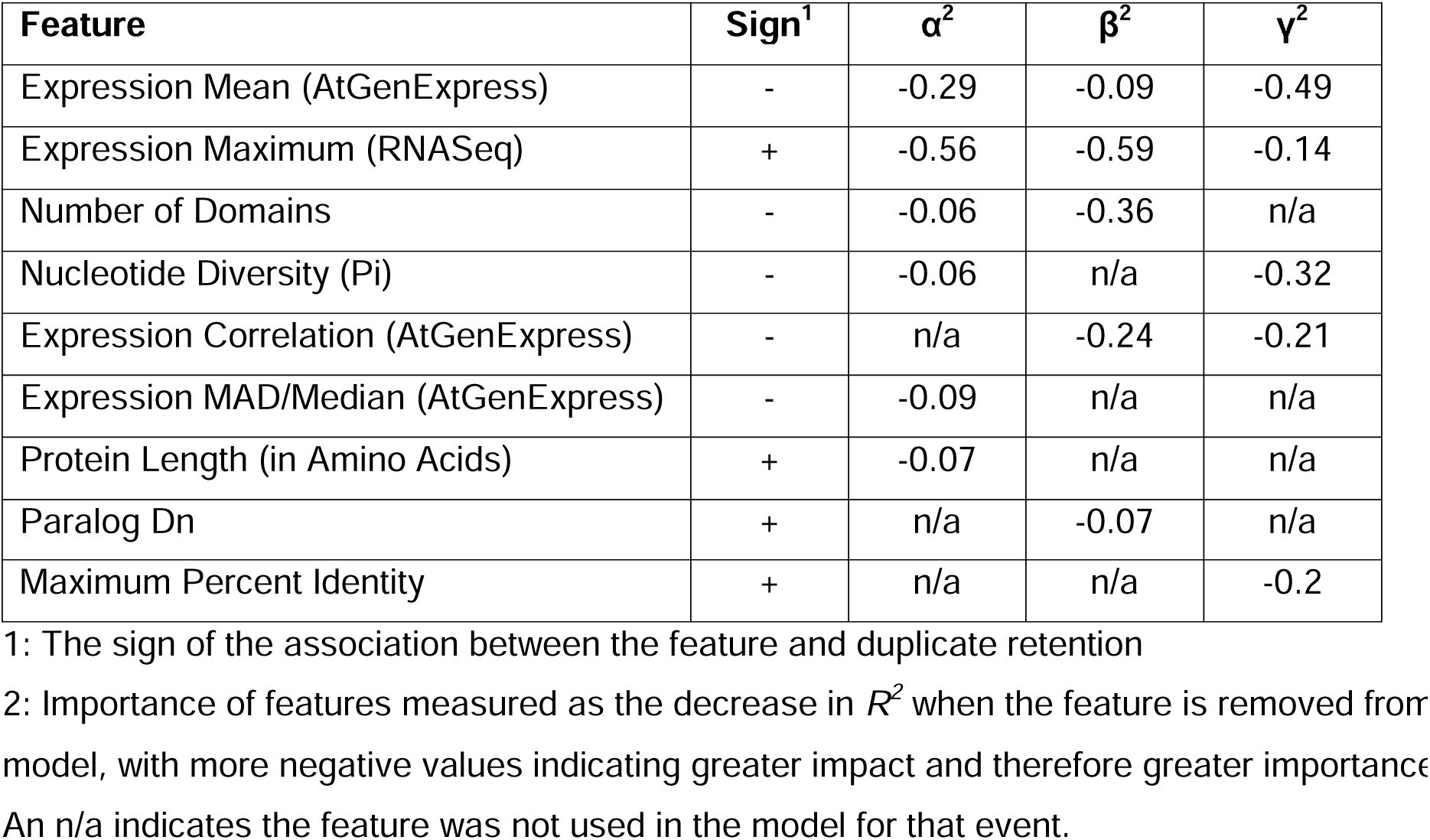
The importance of all features used in the linear models of duplicate retention in function groups across each WGD event

To assess how well the models explained the retention rates of WGD duplicates in different functional groups, we compared the actual and the predicted retention rates. In general, the retention rates predicted by the models closely align with the actual values for each function groups across each event (Fig 2C). However, TF retention rates were consistently underestimated (**Fig S1**), particularly in the γ model the TF retention rate is predicted to be only 76% of the actual value (black arrows, Fig 2C). As such, while our models provide a general explanation for the differences in duplicate retention between function groups, the behavior of TFs departs from the norm. To conclude, we demonstrated that retention rates of genes in different function groups are related to expression level/pattern and sequence divergence. However, while these features are useful for predicting retention rates for some function groups, they systematically underestimated TF retention rates.

### Machine learning classification of duplication status of individual genes

One explanation for the underestimation of TF retention rates is that the relationships between retention rate and feature values of TFs are significantly different than those of other genes, such that a general model gives poor predictions. To explore this possibility, we used a machine learning approach, Random Forest, to classify each individual gene as either having or lacking a duplicate from a WGD event based on the gene’s individual properties (see **Methods**). We constructed separate classifiers for TFs, protein kinases (another function group with high retention but better linear model fit, see white arrows Fig 2C), and all genes regardless of the function groups. For this analysis, duplicates from all three WGD events were combined into one classifier as small numbers of β and γ made the difficult to correctly classify on their own.

To evaluate the performance of our classifiers, we determined receiver operating characteristic curves (ROCs) for each model (Fig 3) and calculated the Area Under Curve (AUC-ROC), a metric that summarizes the ability of the classifier to recover true positive WGD-duplicate genes at different false positive rates. An AUC-ROC of 0.5 indicates that the classifier is no better than randomly labeling genes as having a retained duplicate or not, while an AUC-ROC of 1.0 indicates that the classifier can make predictions without error.

**Figure 3.**
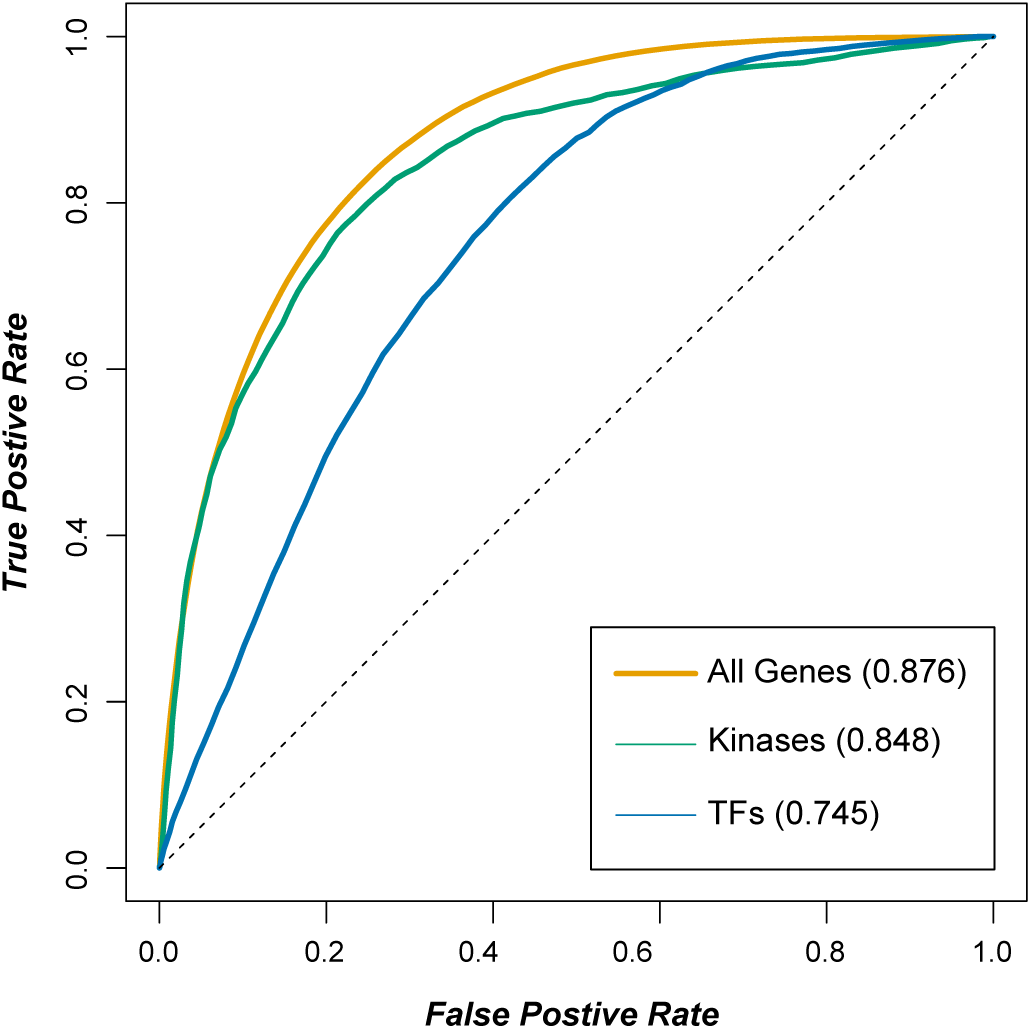
Predicting WGD-duplicate status of *A. thaliana* genes using Random Forest. Receiver-operator characteristic (ROC) curves for Random Forest models predicting duplicate status of all genes in *A. thaliana* (orange), kinases (green), and TFs (blue). The dotted line indicates the curve for random guessing with an area under the ROC curve (AUC-ROC) of 0.5. The AUC-ROC of each model is indicated in parentheses.

Among the classifiers, the one characterizing the full genome performed best (AUC-ROC = 0.86), followed closely by protein kinases (AUC-ROC = 0.82), while the classifier for TFs, while much better than random, did not perform as well (AUC-ROC = 0.74). To investigate the source of the difference, we determined the importance of each feature to the classifier by calculating the Mean Decrease in Accuracy (MDA) which is the average number of genes misclassified across multiple runs as a result of removing a feature (Table 3). Given TFs are the least well predicted, we suspected the informative features for predicting retention in TFs would differ greatly from those for the genome at large and the protein kinases. Contrary to this expectation, the ranking of importance for TF WGD-duplicate prediction was more similar to the ranking of features for the whole genome prediction (Spearman’s rank, ρ = 0.86) than the ranking of features protein kinases to the whole genome prediction (Spearman’s rank, ρ = 0.51). This finding suggests that the feature value distributions of TF WGD-duplicate and WGD-singletons are more similar to the genome at large. Therefore, the reason that TF duplicate prediction model had lower performance was not simply because their feature values were substantially different from other duplicate genes. Instead, the features examined simply have lower importance in general for predicting TF retention (average MDA=11.3) than for other genes (average MDA=47.9), suggesting there are additional features important for TF retention that were not considered. For example, we might expect the number of DNA binding sites to be predictive of duplication status as an indication of the breadth of function of the TF which is related to the probability that a duplicate copy has been retained through subfunctionalization or gene balance.

**Table 3.** The importance (rank) of all features used in the classification of individual duplicate genes

| Feature | Genome<sup>1</sup> | Kinases<sup>1</sup> | TFs<sup>1</sup> |
| --- | --- | --- | --- |
| Maximum Percent Identity (Paralog) | 171.6 (1) | 57.9 (1) | 29.4 (2) |
| Sequence Conservation (Viridiplantae) | 109.3 (2) | 27.2 (2) | 31.8 (1) |
| Gene Family Size (OrthoMCL) | 81.6 (3) | 2.0 (19) | 27.7 (3) |
| Protein Length (in Amino Acids) | 52.5 (4) | 14.2 (3) | 10.5 (11) |
| Expression Breadth (AtGenExpression) | 46.2 (5) | 9.4 (10) | 11.2 (8) |
| Expression MAD/Median (AtGenExpress) | 41.0 (6) | 5.4 (11) | 11.8 (5) |
| Expression Mean (LightDev Data) | 40.8 (7) | 10.4 (7) | 10.7 (9) |
| Expression Breadth (RNASeq) | 40.0 (8) | 3.3 (15) | 11.5 (6) |
| Expression Mean (Control Data) | 39.2 (9) | 10.7 (6) | 10.7 (10) |
| Expression Median (AtGenExpress) | 37.5 (10) | 10.2 (8) | 10.2 (12) |
| Expression Mean (Stress Data) | 37.0 (11) | 11.5 (4) | 12.0 (4) |
| Expression Mean (AtGenExpress) | 36.9 (12) | 11.3 (5) | 11.3 (7) |
| Expression Median (RNASeq) | 34.8 (13) | 4.7 (12) | 4.9 (16) |
| Sequence Conservation (Metazoa) | 34.5 (14) | 3.6 (13) | 4.4 (18) |
| Nucleotide Diversity (Pi) | 32.4 (15) | 9.7 (9) | 6.2 (14) |
| Expression Mean (Diff Data) | 31.6 (16) | 1.7 (2) | 7.9 (13) |
| Sequence Conservation (Fungi) | 30.6 (17) | 2.4 (17) | 4.6 (17) |
| Number of Protein Domains | 28.4 (18) | 2.3 (18) | 5.6 (15) |
| Expression Mean (RNASeq) | 18.6 (19) | 3.0 (16) | 2.9 (19) |
| Expression Maximum (RNASeq) | 12.9 (20) | 3.3 (14) | 0.4 (20) |

1: The importance of the feature as defined by the mean decrease in accuracy of the classification when the feature is removed. Features are ordered according to the rank of their importance in the whole genome model and the rank of each value for each model is indicated by (),

Furthermore, the most informative feature for classifying kinases and the whole genome, the percent identity to the best matching paralog in *A. thaliana,* was less important when applied to TFs (Table 3). Although the maximum percent identity of WGD-duplicates compared to WGD-singletons is significantly higher in full genome (*p* = 1e-320), protein kinases (*p* = 1.1e-36), and TFs (*p* = 6.2e-12), the magnitude of the difference was greater for protein kinases (11.2%) and the whole genome (11.3%) than TFs (4.4%). This is due to WGD duplicate TFs having lower maximum percent identity (71.3%) than either kinases (75.2%, *p*=4.1e-24, t-test) or all genes (72.5%, *p*=5.9e-83, t-test), while WGD-singletons TF had higher identity (66.9%) than kinases (64.0%, p=4.2E-35, t-test) and all genes (61.3%, p=1.9e-223, t-test). This observation may related to non-duplicate TF genes having apparent paralogs more often than non-duplicate genes do on average across the *A. thaliana genome* (**Fig S2**). The variance in the importance of maximum percent identity accounts for most of the performance difference across the classifiers as removing this feature yields similar results from all three (**Fig S3**). Similarly, inflating the difference in the percent identity of TF WGD-duplicates and WGD-singletons from 4.4% to 11.2% (the difference for protein kinases) would raise the predicted retention of TF from the γ WGD from 2.50 to 2.94, making up for more than half of the original error.

We would expect that other features used in our linear models (Table 2) would also be useful for classifying genes within function groups. However, the average importance rank of features found in more than one linear model was low (13.9 of 20), with the maximum expression value in RNA-seq being the worst feature in both the whole genome and TF classifiers. Of the four linear model features, mean expression in AtGenExpress had the highest rank in the whole genome (12th), TF (7th), and kinase classifiers (5th). However, the difference in mean expression between WGD-duplicates and WGD-singletons was not consistent: WGD duplicates genes were more highly expressed across the whole genome (+0.32, p=4.0e-23), and TFs (+0.37, p=1.0e-4), but in protein kinases WGD-singletons were more highly expressed, though not at a significant level (-1.1, p=0.77). Hence, not only does relationship between gene features and retention depend on the gene function, but the relationship within individual function groups can be the opposite direction of the relationship across function groups. For example, the high retention of the TF function group is in part due to relatively low average expression in AtGenExpress, but within TFs, genes with higher average expression are more often WGD duplicates. This suggests that selection for duplicate retention is dependent not only on function and features, but their interaction as well, though the exact nature of these interactions is beyond the scope of this study.

### Partitioning of ancestral expression states following TF duplication

While the gene features (Table 2) were generally useful predictors of WGD-duplicates, they were less useful for predicting TF duplicates specifically. To further explore what characteristics retained TF WGD-duplicates possess, we examined how the functions of retained TF WGD-duplicates have evolved following WGD events. To do this, we first used expression patterns as a proxy of TF function(s) and inferred the likely expression states of the ancestral TFs prior to WGD (see **Methods**). Ancestral expression values were inferred from extant gene expression values that had been discretized into quartiles (expression state = 0, 1, 2, or 3) based on the distribution of expression levels for each array experiment. Additionally, expression data were grouped into four subsets, including control conditions (Ctrl), light and development sets (LightDev), abiotic and biotic stress treatments (Stress), and differential expression between stress treatments and controls (Diff), and analyzed separately. This grouping was used to distinguish between trends that were universal or specific to certain datasets. We were able to infer 165,385 ancestral expression states across 474 TF WGD-duplicate pairs (a detailed breakdown of inferred states can be found in **Table S3**).

First, to test how often the expression states of TFs are retained post-duplication, we compared the expression states of individual, extant TF WGD-duplicate to its inferred ancestral states (Fig 4A). Although all possible changes in expression state were observed between ancestral and extant TFs in each expression data subset, the most common ancestral-extant expression state combination was that the ancestral and extant TFs had the same expression quartiles (diagonal red boxes, Fig 4A). This is true across all expression quartiles, though the deviation from expectation was greatest for expression values in the lowest (0) and highest (3) quartiles. This general pattern holds across all four data subsets (**Fig S4**), suggesting that most TFs WGD-duplicates retain their original expression irrespective of what that expression is. However, when considering a pair of duplicates, we found that when the ancestral state was retained in one duplicate, it was lost more often in the other duplicate than expected by random chance (Fig 4B). This “partitioned” state of TF WGD-duplicates pairs is most over-represented in duplicates from the α event compared to duplicates from the older β and γ events (Fig 4B). In these older WGD events, having neither duplicate inherit the ancestral expression state is more common than the partitioned state where only one copy inherits the ancestral state. Using ANOVA, we confirmed that there is indeed significant interaction between the expression state of a TF WGD-duplicate pair and the timing of the WGD event (*p*<2e-16), which indicated that partitioning occurred relatively quickly after the most recent WGD, but that these partitioned patterns were not necessarily retained as the duplicates age.

**Figure 4.**
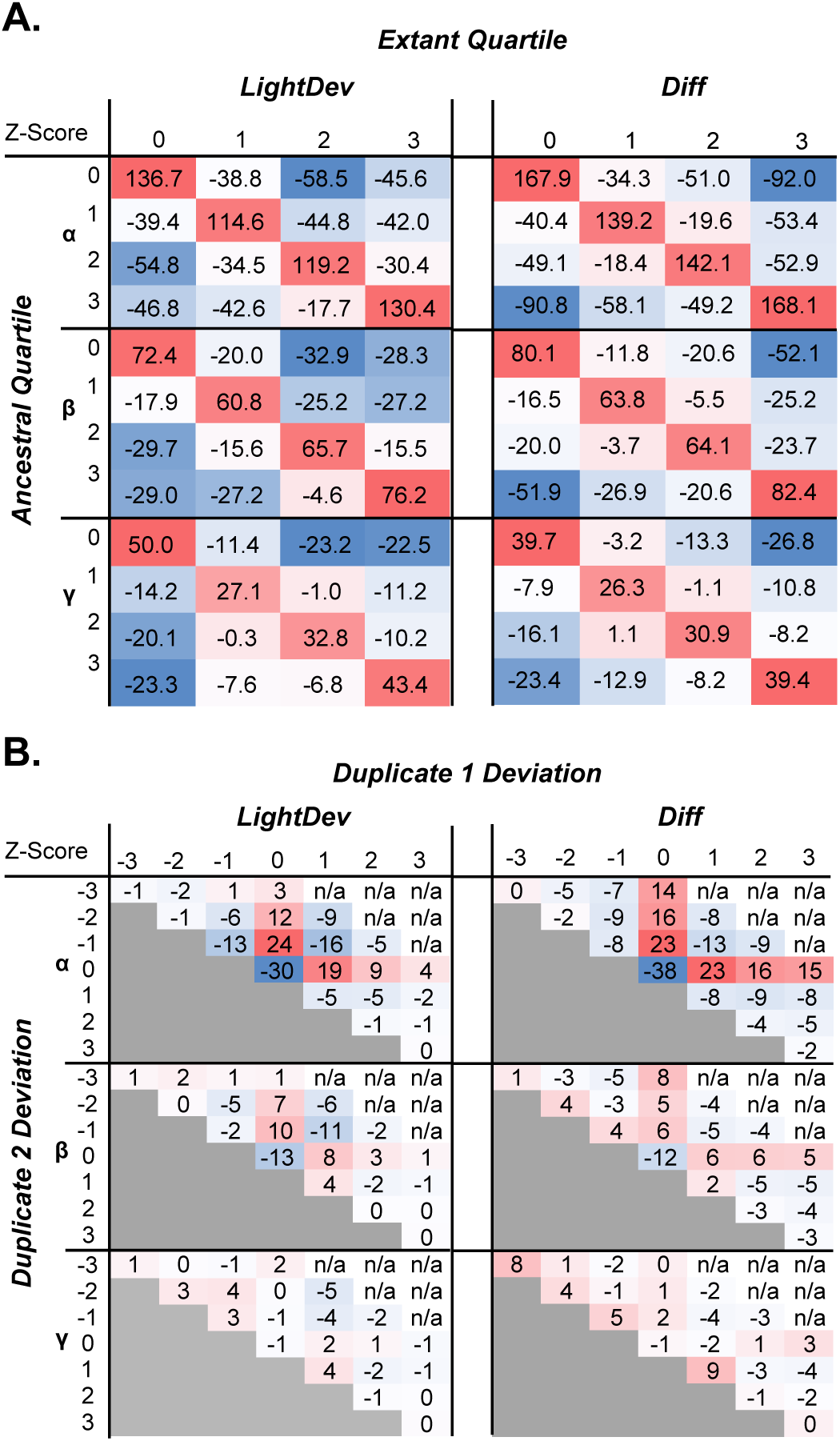
Evolution of expression in TF WGD-duplicates. **(A)** Difference in expression quartile of individual TF WGD-duplicates compared to their ancestral state. Heatmaps show the z-scores of the observed frequency of each difference compared to the expected frequency for LightDev (left column) and Diff (right column) data across all three duplication events (α = top, β = middle, γ = bottom). Color correlates with the magnitude of the z-score, with darker red values indicating counts further above random expectation and darker blue values indicating counts further below random expectation. **(B)** Deviation of pairs of TF WGD-duplicates from their ancestral state, defined as the difference value that each duplicate in a pair has from its ancestral state. Heatmaps show the z-scores of the observed frequency of WGD-duplicate pair deviation compared to the expected frequency for LightDev (left column) and Diff (right column) data across all three duplication events (α = top, β = middle, γ = bottom). Color correlates with the magnitude of the z-score as in (A)

Next we asked if TF duplicates expression tends to increase or decrease when they deviate away from the ancestral state. Because we found a significant interaction between the expressions state evolution of TF WGD-duplicate pairs and the subset of the expression data used (*p*=2.5e-05), we asked this question for each subset of expression datasets individually. For the LightDev (Fig 4B), Ctrl, and Stress expression subsets (**Fig S5**), partitioning of ancestral expression states among duplicates favors small, negative changes from the ancestral states. Based on an earlier study showing that *A. thaliana* duplicates tend to be expressed at a lower level compared to the ancestral state (Zou et al. 2009b), we anticipated TFs would lose their ancestral stress response. However, when we looked at the Diff subset, which looks at differential expression as opposed to raw expression, we found that TFs were equally likely to increase or decrease differential expression in response to stress compared to the ancestral state.

To test the significance of this difference in non-ancestral expression amongst expression subsets, we modeled the evolution of ancestral expression (O) to higher (+) and lower (-) expression states following a WGD duplication event was modeled using ordinary differential equations (see **Methods**). We compared a one-parameter model where the rates of transition from (O) to (+) and (-) were set to be equal to a two-parameter model where the rates to (+) and (-) were allowed to differ (**Fig S6**). The two parameter model was only significantly better than the one parameter model for the LightDev (likelihood ratio test, p=2.2e-11), Ctrl (p=2.7e-3), and Stress (*p*=2.9e-3) subsets. For these subsets, the rate of evolution from (O) to (-) was 1.9~3.1 times more frequent than that from (O) to (+). For the Diff subset, (O) to (-) was only 1.2 times more frequent, which was not significant (*p*=0.43). In summary, these results suggest that the evolution of TF duplicates favors decreasing expression levels relative to the ancestral expression state (Control, LightDev, and Stress). However, when looking at differential expression in response to stress, TF duplicates can evolve in either direction with approximately equal likelihood. Thus, following duplication, TF duplicates may have increased or decreased responses to stress, rather than losing the response altogether.

### Asymmetry in the partitioning of ancestral expression and regulatory sites

Thus far we show that an ancestral expression state tends to be retained by only one copy of a TF WGD-duplicate pair. One outstanding question is whether each copy would retain different parts of the ancestral expression state, as would be expected if the TF duplicates were retained due to subfunctionalization (Force et al. 1999). To address this, we considered all of the partitioned expression states (i.e. all expression series showing partitioning) across a pair of TF WGD-duplicates. If partitioning were random, we would expect that the number of ancestral states retained by a single WGD-duplicate to follow a binomial distribution for the given number of partitioned expression states (n) and a retention probability of 0.5 such that each copy is equally likely to retain ancestral express states. Under this scenario, the expected asymmetry of a duplicate pair (the difference in the fraction of ancestral states inherited between duplicates) is 0.18 given the observed distribution of partitioned states. However, the actual mean asymmetry between TF WGD-duplicates was 0.67, which is unlikely to have been generated by random partitioning (*p*<1e323) expected under the subfunctionalization model. In fact, in 35.1% of cases, one WGD-duplicate retains all the ancestral expression and the distribution is highly biased towards higher asymmetry (Fig 5A). As with the mean asymmetry, the skewed distribution of asymmetry values is also significantly different from what was expected from random partitioning (Kolmogorov-Smirnov test, *p*<2.2e-16). This biased partitioning was also found within the Ctrl (mean=0.84), LightDev (mean=0.67), Stress (mean=0.69), and Diff (mean=0.56) subsets. Given these results, for TF WGD-duplicate pairs we can generally define one duplicate as being “ancestral” and the other as being “non-ancestral”. Why then is the non-ancestral copy being retained? One hypothesis is that the non-ancestral copy is retained because it has acquired a novel function.

**Figure 5.**
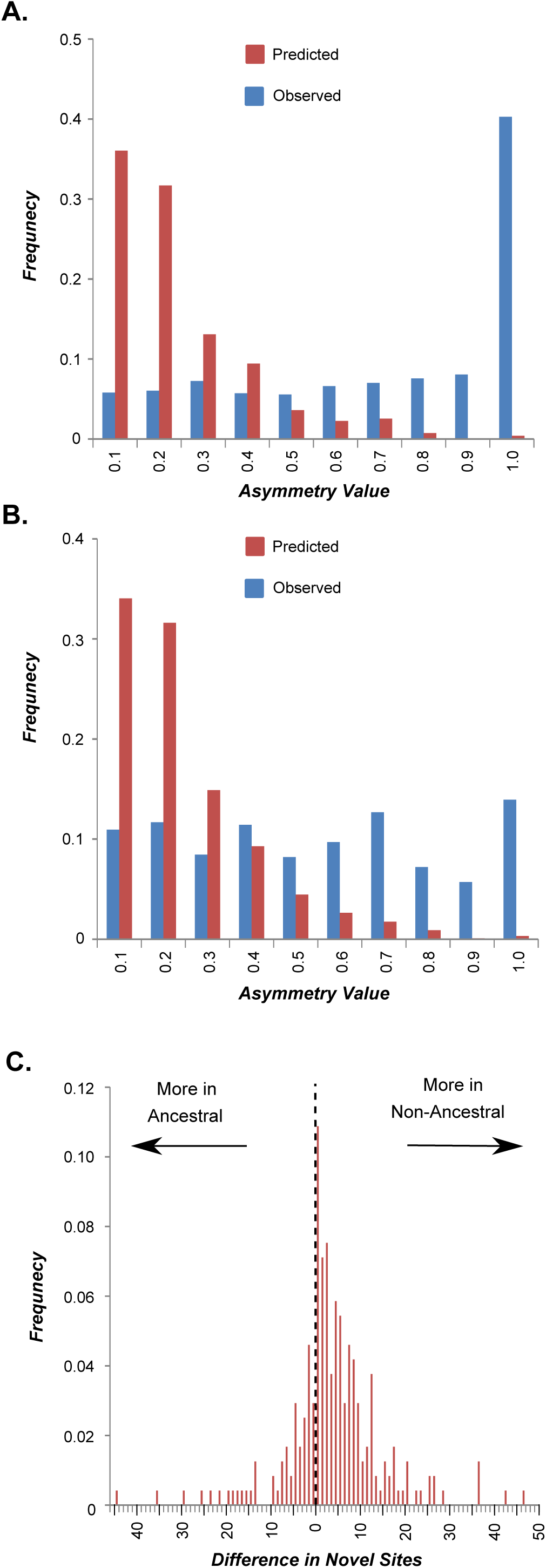
Asymmetry of ancestral state retention in TF WGD-duplicates. (A) The asymmetry value (see Methods) of ancestral expression partitioning between TF WGD-duplicates. (B) The asymmetry values of ancestral *cis*-regulatory site partitioning between TF WGD-duplicates. (C) The frequency distribution of the difference in number of novel *cis*-regulatory sites between ancestral and non-ancestral WGD duplicates. Values on the x-axis to the left of zero indicate the magnitude of differences in the number of sites favoring the ancestral duplicate while values to the right of zero indicate the magnitude of differences favoring the non-ancestral duplicate.

To test whether the non-ancestral copies tend to have novel functions, we first applied our model of ancestral-state partitioning to *cis*-regulatory sites. We used *cis*-regulatory sites here because the discretized expression levels used above allowed us to determine the direction of changes away from the ancestral expression state, but not whether an expression state was novel. The *cis*-regulatory sites used here are from putative binding sites of 345 *A. thaliana* TFs (O’Malley et al. 2016). We applied the same methodology used to infer ancestral gene expression to infer ancestral *cis*-regulatory sites of ancestral TFs (see Methods). Among 16,415 ancestral-extant site comparisons, the majority (58.9%) involved the loss of an ancestral site in only one WGD-duplicate gene, which is significantly different compared what would be expected if WGD-duplicate and ancestral genes were randomly associated (42.3%; t-test, *p*<1e-323). In contrast, retention (15.8%, *p*<1e-323) and loss (10.2%, *p*<1e-323) of ancestral *cis-* regulatory sites in both WGD-duplicates were significantly less frequent than randomly expected. Similar to ancestral expression state evolution, the partitioning patterns of ancestral *cis*-regulatory sites were highly asymmetric (Fig 5B), resulting in a distribution that is significantly different from the distribution generated by random partitioning (Kolmogorov-Smirnov test, *p*< 2.2e-16). Thus, much like what we observed for expression, these results suggest that, with regard to *cis*-regulatory sites, TF WGD-duplicates can be classified into ancestral and non-ancestral copies. Most importantly, amongst the 249 duplicate pairs with at least one novel regulatory site, in 71.0% of cases the non-ancestral copy had more novel *cis*-regulatory sites (Fig 5C). This is significantly different from what would be expected if ancestral site retention and gain of novel regulatory sites associated independently (49.8%, *p* < 3.8e-12). Furthermore, in 61.8% of pairs only the non-ancestral copy had novel sites, while the ancestral copy contained all of the novel sites in only 14.0% of pairs. These patterns suggested that, the gains of novel *cis*-regulatory sites likely contribute to the retention of the non-ancestral TF duplicate copies.

Note that we can divide each pair of WGD-duplicates into an ancestral copy and a non-ancestral copy based on either expression or *cis*-regulatory site information. Given the important roles of *cis*-regulatory sequences in regulating gene expression, we expected that the ancestral and non-ancestral designation defined according to expression data should be similar to that defined based on *cis*-regulatory sites. Among the 179 TF WGD-duplicate pairs with data on gene expression and regulatory sites, 59.8% follow this expected pattern. This percentage was higher than expected by random association (24.6%, *p* = 1.8e-20). After examining the evolution of expression states of TFs and *cis*-regulatory sites controlling TF expression, the next question is how the regulatory targets differed between TF WGD-duplicates. This is currently not feasible because the *A. thaliana* TF-target data set was too sparse for clear inference, however, we have demonstrated that evolution of WGD-duplicates clearly favors the partitioning of ancestral states into distinct ancestral and non-ancestral duplicates.

### Patterns of WGD-duplicate divergences and partitioning results from evolutionary bias

We have demonstrated that partitioning of ancestral expression and regulation into ancestral and non-ancestral duplicates is favored following WGD-duplication of TFs. It remains an open question if this preference is due to bias towards partitioning or simply results from the progressive loss of ancestral function from duplicate genes. One possible explanation for the observed frequency of partitioning is that it results from the timing of WGD duplications in the *A. thaliana* lineage. Assuming that mutations occur and are fixed by drift, the time required for mutations that alter expression patterns to accumulate in two WGD-duplicated loci is expected to be longer than that for a single locus. Under the scenario where all mutations occur and are fixed at a constant rate, there would be a time when we would expect to find partitioning of ancestral state enriched simply because duplicate pairs are more likely to have lost the ancestral state at a single locus than both loci or neither loci. In contrast, if there is bias for the partitioning of ancestral states, we would expect the loss of the ancestral state at the first locus of a TF WGD-duplicate pair to occur must faster than at the second locus.

To determine whether the observed frequency of partitioning of ancestral states requires biased evolution or not, we modeled loss of ancestral state at pairs of loci in TF WGD-duplicates using a system of ordinary differential equations (see **Methods**). Using the synonymous substitution rate of TF WGD-duplicate pairs derived from the α, β, or γ events as a proxy for time, the rate of transition between WGD-duplicate pairs where both (state II), only one (state I) and neither (state O) duplicate had lost ancestral expression was modeled (Fig 6A). We compared a model where the rates of transitions between all states were equivalent (one-parameter model, Fig 6B) with a model where the transition rates between state II and I were allowed to vary from those between state I and O (two-parameter model) (Fig 6C). These models were applied to all expression subsets, the results for the LightDev dataset are shown in Fig 6 and the remainder can be found in the supplement (**Fig S7**).

We found the two-rate model to be significantly better at explaining the observed difference in WGD-duplicate states over time (Likelihood Ratio Test, p-value < 2e-14). Regardless of the expression data set, the transition rates between state O (ancestral expression in both duplicates) and I (ancestral expression in on duplicate) were 7-13 times higher than the rates between state I and II (ancestral expression in neither duplicate). Given that a pair of TF WGD duplicate would have the same expression patterns as their ancestral gene initially (state O), our finding suggests that the number of partitioned WGD-duplicates accumulated relatively rapidly post WGD, followed by slow accumulation of WGD-duplicates pairs where ancestral expression had been lost entirely. Applying this same approach to model regulatory site evolution revealed an even more extreme difference, as the rates governing the transition between state O and I are two orders of magnitude higher than between state I and II (Fig 6E-G). Additionally, since the best fit model for the regulatory data involved allowing all four rate parameters to vary (p-values 4.8e-13 and 1.2e-11 vs. 1 and 2 parameter models respectively), we can state that rates for transition to state I are higher than the accompanying transition rates away (Fig 6H). This pattern does not hold when the four parameter model is fit to ancestral expression state (Fig 6D), however, the model overall was not significantly better than the two parameter model at a threshold of p-value < 0.05, implying that faster transition between O and I compared to I and II is a better at explaining the evolution of expression states.

**Figure 6.**
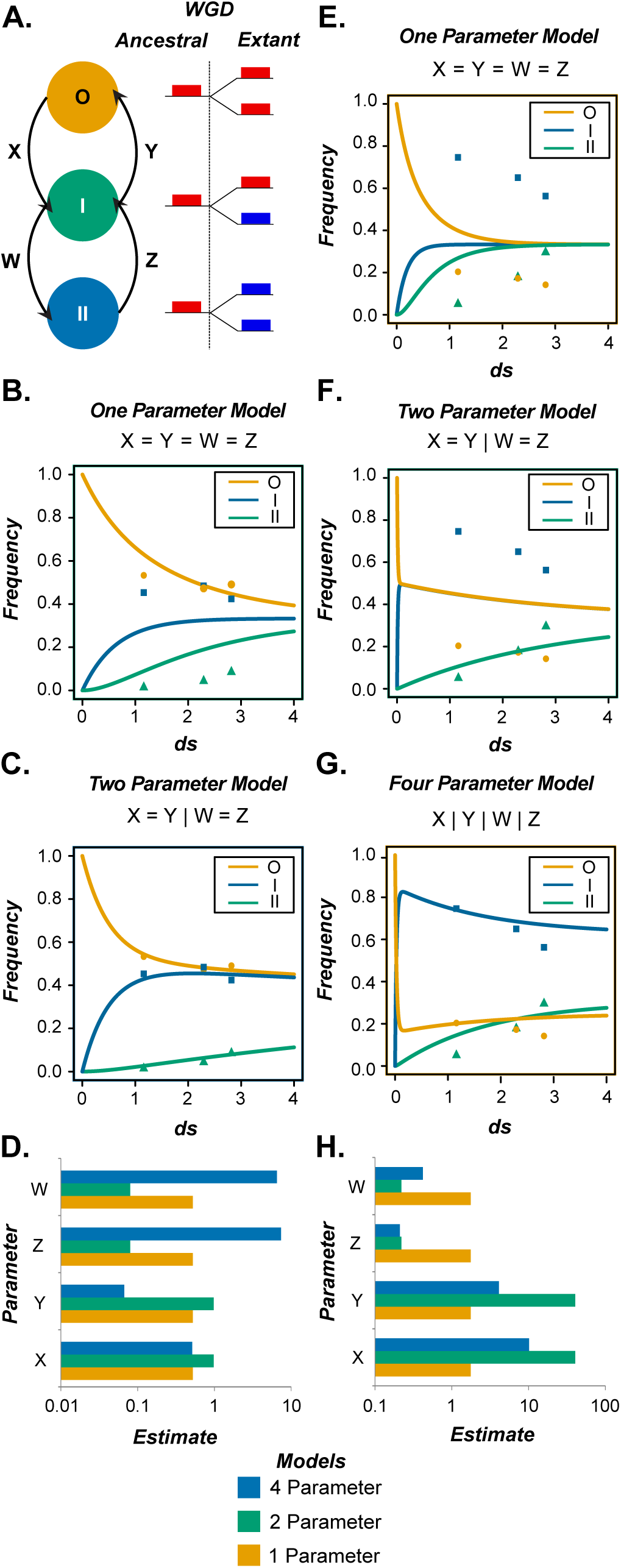
ODE models of TF WGD-duplicate expression and *cis*-regulatory site evolution relative to the ancestral state **(A)** In this model, we consider the transition of WGD-duplicate pair expression between three possible states relative to their ancestral state (O = both retained, I = one retained, II = neither retained) using four variables representing the rate of transition between state (x,y,w,z). **(B)** Results for the one parameter version of the model (z=y=w=z) showing the change in time (x-axis) of the frequency (y-axis) of each WGD-duplicate-pair state (O = orange, I = blue, II = green). Curves represent the continuous output of the model with symbols indicate the observed values on which the models were built (O = circle, I = square, II = triangle). **(C)** Results for the two parameter version of the model (z=y|w=z). Axis, color, lines and symbols are used the same as in (B). **(D)** Bar graph of the parameter values (x,y,w,z) for the one (orange), two (green), and four (blue) parameter versions for the expression ODE model. **(E)** Results for the one parameter version of the model (z=y=w=z) showing the change in time (x-axis) of the frequency (y-axis) of each WGD-duplicate-pair state (O = orange, I = blue, II = green). Curves represent the continuous output of the model with symbols indicate the observed values on which the models were built (O = circle, I = square, II = triangle). **(F)** Results for the two parameter version of the model (z=y|w=z). Axis, color, lines and symbols are used the same as in (E). **(G)** Results for the two parameter version of the model (z|y|w|z). Axis, color, lines and symbols are used the same as in (E). **(H)** Bar graph of the parameter values (x,y,w,z) for the one (orange), two (green), and four (blue) parameter versions for the *cis*-regulatory site ODE model.

The non-equivalency in the rate at which the ancestral state is lost in the first and second duplicates indicates that the frequency of partitioning is not trivial, but rather results from a bias against losing the ancestral state from the second TF WGD-duplicate relative to the first. Furthermore, in the case of ancestral regulatory sites, we found that evolving towards a partitioned state was always favored over the corresponding path away from partitioning. This suggests that partitioned TF WGD-duplicates pairs do not accumulate simply because of selection against having both duplicates ancestrally expressed, but that maintaining the original number of ancestral states is favored. In summary, these results combined with the finding that novel *cis*-regulatory sites tend to accumulate in the non-ancestral duplicate, suggest that partitioned WGD-duplicate TFs result from selection on one copy that has neofunctionalized while the other maintains the ancestral gene function.

## Discussion

WGD is unique amongst the mechanisms of gene duplication in that all loci are duplicated and thus affected equally. As such, any bias in duplicate retention must occur in the aftermath of the WGD event as the genome experiences further rearrangement and reduction. In allopolyploids, where WGD is the consequence of merging two related parental genomes, the process of fractionation (i.e. gene losses after WGD) is biased with regards to the parent of origin and the distribution of deleterious mutations (Thomas et al. 2006; van Hoek and Hogeweg 2007; Schnable et al. 2011). However, we would not expect this to result in different retention rates amongst function groups unless genes with certain functions have different predispositions to loss of function following WGD. Therefore, the significant differences in the frequency of duplicate retention between function groups are likely the result of selective pressures on these gene functions relative to their duplicate status.

It is well established that genes with certain molecular functions are enriched among retained WGD-duplicates (Blanc and Wolfe 2004; Carretero-Paulet and Fares 2012), including protein kinases and TFs (Maere et al. 2005; Shiu et al. 2005). However, the reason these genes are enriched amongst WGD-duplicate was unknown. In this study, we have shown that duplicates are retained at different rates across function groups depending on their expression, conservation, and structure, suggesting these features are related to the selection for or against retaining a duplicate pair. Importantly, the trends that apply to function groups in general do not necessarily hold for individual duplicates, suggesting unaccounted for interaction between function and features. In addition, the retention rate of TF WGD duplicates, once normalized against the retention rates of other genes in the genome, is higher for older duplication events.

We propose two explanations for this observation: (1) TF duplicates produced by the γ event were retained more frequently than duplicates produced by later events, (2) TF duplicates retained from the γ event were more frequently retained following subsequent duplication by later events (see **Supplemental File S1**). It has been shown that duplicate retention in *A. thaliana* is affected by whether the ancestral copy had been transposed prior to the WGD event (Woodhouse et al. 2011), so the notion of prior genome duplicate/rearrange events effecting subsequent retention is not unprecedented. Distinguishing these two scenarios would provide insight into how subsequent duplication events interact with existing WGD duplicates, but in either case, our results suggest non-equivalence either between WGD events or between WGD-duplicates based on their duplication state prior to a WGD event.

Why is there an apparent bias in favor of retaining TF WGD-duplicates? Surprisingly, despite the fact that the TF duplicates were generated ~50 and ~140 million years ago for the α and β WGD events, respectively; retention of ancestral expression is still the most common outcome and even amongst our 1,239 partitioned duplicate pairs, 83.1% have retained ancestral expression in both copies under ≥ 5 other conditions. The frequency with which ancestral expression is conserved suggests that the retention of a significant number of TF duplicates is not due to selection acting on their divergent functions. Furthermore, the fact that a single duplicate pair can both retain ancestral expression under some conditions and be non-randomly partitioned under other conditions, raises the possibility that both subfunctionalization (Force et al. 1999) and gene balance played a role in retention of these WGD-duplicates (Birchler and Veitia 2007; Birchler and Veitia 2010). The question then becomes whether initial retention by gene balance creates the opportunity for later subfunctionalization or if gene balance restrains the divergence of ancestral expression in duplicates pairs that have already been selected for.

The observed evolution of extant gene expression post duplication demonstrates a preference for maintaining partitioned functional states amongst retained TF WGD-duplicate pairs where one duplicate mirrors the ancestral gene’s functional states while the other copy inherits significantly fewer ancestral states. On an experiment-by-experiment basis, the partitioning of ancestral expression states appears to support the notion of WGD-duplicate retention by subfunctionalization (Force et al., 1999). However, when examining the ancestral state partitioning patterns across multiple experiments, we find an extreme bias where one TF retains most of the ancestral states and other, non-ancestral, copy retains few. Most importantly, the non-ancestral copy tends to gain novel *cis*-regulatory sites. This pattern harkens back to the notion of there being an ancestral copy and a neofunctionalized copy after duplication, contributing to the retention of both duplicates (Ohno, 1970). Thus, for duplicate copies with significant expression and *cis*-regulatory site differences, both neo-functionalization and subfunctionalization are likely important for duplicate retention, with the former playing a more important role.

Rather than suggesting a singular explanation of why TF WGD-duplicates are retained, our findings suggest a nuanced pattern of expression and *cis*-regulatory state evolution between duplicates. This raises the question of what are the relative contributions of theorized models of duplicate retention, such as subfunctionalization, gene balance, neo-functionalization, and escape from adaptive conflict. One way to assess contributions of different retention models would be to extend our differential equation models to include more recent WGD events. The a, β, and γ events are relatively ancient taking place ~50-300 million years ago (Bowers et al. 2003). As such, any part of our model prior to the α event, including the initial rapid drop of WGD-duplicates in state O, are extrapolated from what best fits the data from the α, β, and γ events. In addition, for the most ancient γ event, there are so few duplicates that the model quality is significantly impacted. Thus, instead of focusing on even more ancient events, incorporating earlier events could reliably resolve the initial behavior of WGD-duplicate and elucidate the roles of different models of retention. For example, were data from more recent WGD events to indicate initial lag period prior to the accumulation of WGD-duplicates with partitioned ancestral functions, it would support the hypothesis that future sub-and/or neo-functionalization is enabled by initial retention because of gene balance (Veitia et al. 2013).

Additionally, while WGD-Duplicates TFs are known to be preferentially retained across many plant species (Carretero-Paulet and Fares 2012), the patterns of ancestral expression and regulatory site partitioning we have uncovered is in *A. thaliana.* It remains unclear if these patterns are specific to *A. thaliana,* shared by other plant lineages sharing the α, β, and γ events, or is common to any species with similar patterns of WGD events. Even if the overall pattern of TF evolution is consistent across multiple species, it is possible that TF families may evolve differently from each other. In future studies, it will be important to directly compare the size, rate of retention, and rate of partitioning both within and across species in individual families. A multi-species comparison will also be informative for further our understanding of the interplay between WGD and TF retention. For example, we found that γ duplicate TFs have a relatively higher rate of retention compared to the α and β events. Comparing retention of duplicate TFs across multiple species which share the γ duplication event, but have independent subsequent WGDs would allow us to assess how older and younger WGD events may jointly influence TF retention.

Although there are many questions yet to be answered about the factors affecting whether or not a duplicate pair is retained following WGD, we have found that the frequency with which duplicate genes are retained following a WGD events is strongly correlated with the expression, conservation, and structural features of that gene. And although our general model does not give perfect predictions for all function groups, it can serve as a basis for exploring more complicated interactions underlying duplicate retention, namely, the potential interaction between gene features and annotated gene function suggested by our results. Furthermore, the partitioning of ancestral expression and the non-random distribution of ancestral and new *cis-* regulatory sites suggest that selection for both existing and novel functions plays a major role in the retention of TF duplicates. In contrast, most duplicates retain ancestral expression levels under some conditions in both duplicates, making it likely that the complete retention of ancestral expression is preferred under certain conditions. Further investigation of which genes and, more specifically, which expression/regulatory states are preferentially retained, partitioned, or neofunctionalized will benefit from the modeling approaches developed here and increase our knowledge of how duplication interacts with gene function

## Methods

### Genome sequence, gene annotation, and Expression Data

Genome sequences, protein sequences, and gene annotation information for *A. thaliana* was obtained from Phytozome v10 (https://phytozome.jgi.doe.gov/pz/portal.html). WGDs were defined according to Bowers et al. (2003) and tandem genes in *A. thaliana* were defined as pairs of reciprocal best BLAST hits with an e-value < 1e-10 and ≤ 5 intervening genes. Expression microarray data for this study was taken from AtGenExpress (Schmid et al. 2005; Kilian et al. 2007; Goda et al. 2008), normalized using RMA (Irizarry et al. 2003) in R as performed previously (Zou et al. 2009a), and divided into four groups: control conditions (in environmental condition experiments, Ctrl), light and development set (LightDev), abiotic and biotic stress treatments (Stress), and differential expression between stress treatments and controls (Diff). The individual experiments in the Controls, Development and Light, and Stress Treatment groups are described in **Table S4**. The Diff data contains the log2 normalized difference between data sets for each stress condition/treatment/duration and its corresponding controls. In addition to microarray data, we have included a set of 214 RNA-sequencing samples (**Table S5**) from *A. thaliana* Col1 wildtype from the Sequence Read Archive (https://www.ncbi.nlm.nih.gov/sra). Raw sequence reads were processed using Trimmomatic (Bolger et al. 2014), with a quality threshold of 20, window size of 4, and hard-clipping length of 3 for leading and trailing bases. Processed reads were then mapped to the *A. thaliana* genome using Tophat2 (Kim et al. 2013) and expression levels calculated with Cufflinks (Trapnell et al. 2010), both with a maximum intron length of 5,000bp.

### Defining TFs and other groups of genes in *A. thaliana*

TFs were defined according to the criteria used by the Plant Transcription Factor Database (Jin et al. 2014), which has annotated 1,717 unique TF loci in *A. thaliana.* Additionally, 19 additional function groups were defined using Gene Ontology (GO) Terms in the molecular function and biological process categories (The *Arabidopsis* Information Resource, https://www.arabidopsis.org/), each containing 100-2,000 genes and ≥20 WGD-duplicate pairs. We excluded GO:0006355 (regulation_of_transcription, DNA-templated) due to its substantial overlap with the TF group we have defined above and because this category include other types of regulators in addition to DNA-binding TFs. The GO terms for the other function groups include: ATP Binding (GO:0005524), catalytic activity (GO:0003824), defense response (GO:0006952), DNA endoreduplication (GO:0042023), hydrolase activity hydrolyzing O-glycosyl compounds (GO:0004553), kinase activity (GO:0016301), lipid binding (GO:0008289), oxidoreductase activity (GO:001649), oxygen binding (GO:0019825), protein binding (GO:0005515), proteolysis (GO:0006508), response to auxin (GO:0009733), response to chitin (GO:0010200), RNA binding (GO:0003723), transferase activity, transferring glycosyl groups (GO:0016757), translation (GO:0006412), transporter activity (GO:0005215), ubiquitin-protein transferase activity (GO:0004842), zinc ion binding (GO:0008270). A list of genes in each group can be found in **Table S6**.

### Fitting duplicate retention rate within each group of genes for each WGD event using linear models

A gene was designated as a WGD-duplicate if its paralog derived from a particular WGD event is present. For a gene without its paralog from WGD, it was designated as a WGD-singleton gene. The retention rate for each function group, *g*, after a specific WGD event, w, is defined as:

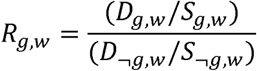

Where *D¬_g,w_* and *D_gw_* are the numbers of WGD-duplicate genes in group *g* and those not in group g (¬g), respectively. *S¬_g,w_* and *S_g,w_* are the numbers of WGD-<u>s</u>ingleton genes in group *g* and those not in group g (¬g), respectively. For each gene group/WGD event, we established a general linear model with the glm function in the R environment which relates the *R_g,w_* to a set of features of each gene group. The 34 features (predictor variables) (**Table S2**) were filtered to prevent over-fitting because the number of observed retention rates was 20. We calculated the correlation between all features to find all cases where the absolute value of correlation was > 0.7. The considerations for which features to keep included: (1) how well each feature correlated with *R_g,w_* on its own, (2) whether the feature was derived from a subset of another feature, and (3) the number of other features with a correlation > 0.7 (favored the elimination of more features). In addition to the above criteria, one data set (protein-protein interactions) was eliminated because of a high frequency of missing values (88%). The synonymous substitution rate (*d_S_*) feature and any feature using *d_S_* in their calculation were also excluded because they would be highly correlated with WGD timing and confound our analyses comparing the three WGD events. The filtering step left 11 features for building the general linear model in the following, iterative fashion. Following fitting the glm function, features were ranked according to their *p* values from the least to the greatest and the feature with the largest *p* value was dropped. The model was then fit to the reduced feature set and features were once again ranked. This process was repeated until the F-statistic (a measure of goodness of fit of the given model against a null model where all coefficients are set to zero) of the model was maximized and the final *p* value was calculated based on the maximal F-stat. The final model for each event can be found in **Supplementary File S2**.

### Predicting WGD-duplicate retention status of individual genes using machine learning

In addition to the linear model, machine learning models for each group of genes (TFs, kinases, or all genes in the genome) were generated to predict whether a gene in a particular group had a retained WGD paralog from either the α, β, and γ event. The machine learning was performed using the Random Forest algorithm implement in the R package randomForest (https://cran.r-project.org/web/packages/randomForest/index.html). We filtered the gene level feature set from a previous study (Lloyd et al. 2015) by removing those with missing values for ≥5% of genes. For the remaining features, missing values were imputed with the rfImpute algorithm in randomForest using 10 iterations of 500 trees. The final matrix of genes and features for TFs, kinases, and the whole genome can be found in **Tables S7, S8,** and **S9,** respectively. Using the imputed data set for each group of genes and for each WGD event, we ran the Random Forest algorithm 10 times with 500 trees (each time with 10 fold cross validation) and collected the resulting votes (retained or not) for constructing Receiver Operating Characteristic curves (ROCs). The importance of each individual feature was assessed using Mean Decrease in Accuracy (MDA), the average number of genes misclassified across multiple runs as a result of removing the feature in question. The statistical significance of the difference in values of a feature between WGD-duplicates and WGD-singletons was determined using Welch’s t-test.

### Inferring ancestral expression levels and *cis*-regulatory sites

DNA-binding domains were identified in TF protein coding sequences using hmmscan via HMMER3 (Mistry et al. 2013) based on the Pfam-A HMMs (version 29.0, Finn et al. 2016) with a threshold e-value of 1e-5. TFs were classified into families according to their DNA-binding domains and 44 of 59 TF families with ≥4 members were used for further analysis (**Table S10**). For each TF family, full-length protein sequences were aligned using MAFFT (Katoh and Standley 2013) with default parameters. The phylogeny of each TF family was obtained using RAxML (Stamatakis 2014) with the following approach: rapid Bootstrapping algorithm, 100 runs, GAMMA rate heterogeneity, and the JTT amino-acid substitution model. These trees were then mid-point rooted with retree in PHYLIP (Felsenstein, 1989). These trees were then used to infer the ancestral gene expression states and the *cis*-regulatory sites of WGD-duplicate TF pairs with BayesTrait (Pagel et al. 2004) as was done in our earlier study (Zou et al. 2009a). The expression data sets used are described in **Table S4**. The discretized gene expression state (0,1,2,3) was based on the quartiles of gene expression levels within each experiment. Thus the inferred, ancestral expression state was also discretized. For *cis*-regulatory sites the binding targets of 345 *A. thaliana* TFs were defined based DAP-Seq data generated by O’Malley et al. (2016). All in-vitro binding targets from the Plant Cistrome Database

(http://neomorph.salk.edu/dap_web/pages/index.php) where 5% of the read associated with a site were found to be in the 200bp peak region. The inference was whether a particular site was present or absent (0,1). For both expression and DAP-Seq data, in cases where there was a missing value, it was explicitly included as an ambiguous state. To call the ancestral state from the expression or *cis*-regulatory site data, we required a posterior probability >0.5. Cases where the called state was ambiguous or no majority existed were excluded from further analysis.

### Asymmetry of the retention of ancestral expression and regulatory sites

For determining expression state asymmetry, only TF WGD-duplicates with ≥5 partitioned ancestral expression states in one of the four expression datasets (Ctrl, LightDev, Stress, and Diff) were considered. For a WGD-duplicate pair with genes A and B, if the number of inherited ancestral expression states in A was larger or equal to that in B, then A and B were defined as the ancestral and the non-ancestral duplicate copies, respectively. The degree of asymmetry (Y_A,B_) of expression states between two duplicates was defined as:

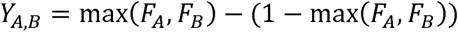

Where F_A_ and F_B_ are the frequency with which ancestral expression was retained across the partitioned states for duplicates A and B respectively. By definition, F_A_ + F_B_ = 1, such that Y_A,B_ is value between 0 and 1 which equals 0 when F_A_ = F_B_ (no asymmetry) and 1 when either F_A_ or F_B_ = 1 (maximum asymmetry).

With the asymmetry values for each TF pair, an average asymmetry value was calculated for each expression dataset, as well as for the union of all WGD-duplicates from all datasets (1,239 values total) to assess how the observed degree of asymmetry compared to what would be expected from random partitioning. The expected distribution of asymmetry values for the expression states of TF WGD-duplicates was determined by randomly partitioning a data set with the same number of pairs and the same distribution of expression states amongst pairs 1,000 times.

For *cis*-regulatory site asymmetry, only TF WGD-duplicates with ≥5 inferred ancestral *cis*-regulatory sites we considered (402 WGD-duplicate pairs total). Similar to expression state asymmetry, in each duplicate pair the ancestral and non-ancestral duplicates were defined according to the number of inherited ancestral sites. For each WGD-duplicate pair, the degree of asymmetry of *cis*-regulatory site among a TF pair was defined analogous to what was done for expression. The expected distribution of asymmetry values for the *cis*-regulatory sites of TF WGD-duplicates was determined by randomly partitioning a data set with the same number of pairs and the same distribution of expression states amongst pairs 1,000 times

### Ordinary differential equation models of TF function evolution

The change in expression states from the ancestral expression quartile to either a higher or lower quartile in an extant TF was modeled as a system of ordinary differential equations such that:

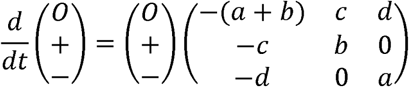

Where *O*, +, and - are the frequency of TF WGD duplicate genes retaining the ancestral expression states, a higher than ancestral expression, and a lower than ancestral expression, respectively. The parameters a, b, c, and d, define the transition rate between these states. This system of equations was solved in Maxima (http://maxima.sourceforge.net/index.html) and best parameters for the observed distribution of duplicates pairs were determined using maximum likelihood estimates calculated with the bbmle package in R (https://cran.r-project.org/web/packages/bbmle/index.html). Non-linear minimization was used to approximate an initial guess, although the actual initial parameters often needed to be adjusted to reach a convergent solution. The best fit parameters for this single duplicate expression state evolution model can be found in **Table S11.**

The loss of ancestral expression states in a pair of duplicated TFs was modeled as a system of ordinary differential equations such that:

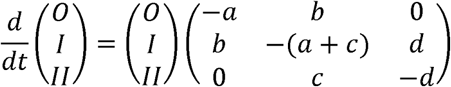

Where *O, I,* and *II* are the frequency of TF WGD duplicate pairs where both, one, or neither duplicate retained the ancestral expression state. The parameters a, b, c, and d, define the transition rate between these states. This system of equations was solved and the initial and best parameters were estimated in the same fashion as above. The best fit parameters for this pairwise expression state evolution model can be found in **Table S12**. The same model was also applied to ancestral regulatory sites with *O, I,* and *II* representing the frequency of TF WGD duplicate pairs where both, one, or neither duplicate retained the ancestral regulatory site.

## Acknowledgements

We thank Johnny Lloyd and Zing Tsung-Yeh Tasi for their advice regarding Random Forest setup and analyzing the importance of predictive features. This work was in part supported by National Science Foundation IOS-1546617 and DEB-1655386 to S.-H.S.; an MSU Discretionary Funding Initiative grant to S.-H.S.

